# Genomics and resistance assays inform the management of two tree species being devastated by the invasive myrtle rust pathogen

**DOI:** 10.1101/2024.10.30.612564

**Authors:** Stephanie H Chen, Jia-Yee S Yap, Veronica Viler, Craig Stehn, Karanjeet S Sandhu, Julie Percival, Geoff S Pegg, Tracey Menzies, Ashley Jones, Karina Guo, Fiona R Giblin, Joel Cohen, Richard J Edwards, Maurizio Rossetto, Jason G Bragg

## Abstract

Myrtle rust is a plant disease caused by the invasive fungal pathogen *Austropuccinia psidii* (G. Winter) Beenken, which has a global host list of 480 species. It was detected in Australia in 2010 and has caused the rapid decline of native Myrtaceae species, including rainforest trees *Rhodamnia rubescens* (Benth.) Miq. (scrub turpentine) and *Rhodomyrtus psidioides* (G.Don) Benth. (native guava). *Ex situ* collections of these species have been established, with the goal of preserving remaining genetic variation. Analysis of reduced representation sequencing (DArTseq; *n* = 444 for *R*. *rubescens* and *n* = 301 for *R*. *psidioides*) showed genetic diversity is distributed along a latitudinal gradient across the range of each species. A panel of samples of each species (*n* = 27 for *R*. *rubescens* and *n* = 37 for *R*. *psidioides*) were resequenced at genome scale, revealing large historical effective population sizes, and little variation among individuals in inferred levels of deleterious load. In *Rhodamnia rubescens*, experimental assays (*n* = 297) identified individuals that are putatively resistant to myrtle rust. This highlights two important points: there are tangible pathways to recovery for species that are highly susceptible to rust via a genetically informed breeding program, and there is a critical need to act quickly before more standing diversity is lost.

## Introduction

The incursion and spread of pests and diseases disrupts natural ecosystems and rapidly alters the population trajectory of native species. This is a key concern in conservation biology (Vitousek et al., 1996) as invasives can ultimately lead to species extinction (Mooney & Cleland, 2001). It is estimated that 65–85% of all plant pathogens are alien species (Pimentel, 1993) and exotic pathogenic fungi proliferate through novel host-pathogen interactions and are often underrepresented in invasion ecology (Desprez-Loustau et al., 2007). A range of coordinated conservation actions are needed to curtail the loss of biodiversity resulting from these introduced plant pathogens (Langhammer et al., 2024).

*Austropuccinia psidii* (G.Winter) Beenken is an obligately biotrophic fungal pathogen that causes myrtle rust. The disease results in deformed leaves, heavy defoliation of branches, reduced fertility, dieback, stunted growth, and even death of mature plants (Carnegie & Pegg, 2018). The pathogen is native to forests of Central and South America, and has expanded its range to the Caribbean, United States, Japan, Australia, China, South Africa, New Caledonia, Indonesia, Singapore, and New Zealand (Chock, 2020). Globally, over 480 species in the plant family Myrtaceae are known hosts (Soewarto et al., 2019).

The impact of myrtle rust diseases is of particular concern to biodiversity in Australia where flora from Myrtaceae dominates (c. 70 genera and c. 1,700 species). The myrtle rust causing pathogen was first detected in Australia in 2010 on the central coast of New South Wales (Carnegie et al., 2010). It has been observed to infect 392 Australian taxa (Soewarto et al., 2019). To date, only the pandemic biotype is present in Australia (da S. Machado et al., 2015). Rust spores have spread and infected host plants across eastern Australia (Carnegie et al., 2016) — from far north Queensland down to Victoria (Victorian Government Department of Energy, Environment and Climate Action, 2023) and Tasmania (Tasmanian Government Department of Natural Resources and Environment, 2024) — and to the Tiwi Islands in the Northern Territory (Westaway, 2016), and recently in 2022 to Western Australia (Government of Western Australia Department of Primary Industries and Regional Development, 2022). No effective control is currently available to protect susceptible species in the wild as spores are readily dispersed via wind, water, and animals. However, for isolated areas such as Lord Howe Island, early detection and rapid response can feasibly eradicate myrtle rust. Therefore, on-ground action is crucial to secure the future of myrtle rust impacted species. Myrtle rust has caused extensive dieback in a range of native taxa, with those most affected typically from coastal heath, wetland and rainforest habitats (Makinson, 2018).

Two previously widespread myrtaceous rainforest species, *Rhodamnia rubescens* (Benth.) Miq. and *Rhodomyrtus psidioides* (G.Don) Benth. are of particular concern because of their high susceptibility to rust (Carnegie & Pegg, 2018; Fernandez Winzer et al., 2019). Prior to the spread of rust, these rainforest species were common in coastal districts and escarpment ranges along the east of Australia. *Rhodamnia rubescens*, or scrub turpentine, is a tree that grows up to 25 m. *Rhodomyrtus psidioides*, or native guava, is a shrub or small tree that grows up to 12 m. Both species have suffered widespread losses throughout their distributions, as exemplified by high mortality rates across surveyed populations in *R*. *psidioides* two to three years following the establishment of *A. psidii* (Carnegie et al., 2016). It has been predicted that 16 rainforest tree species, including *R*. *rubescens* and *R*. *psidioides,* may become extinct in the wild within a generation (Fensham et al., 2021). Given their rapid decline, both species were listed as critically endangered under relevant state and federal legislation.

To prevent the extinction of these species, cuttings have been collected across their ranges, grown as *ex situ* insurance populations in nurseries, and protected with fungicide (Makinson et al., 2020). It is important that we characterise the genetic variation in these collections, and across remaining natural populations, to guide their management. Small populations face several important genetic risks. Loss of genetic diversity can reduce the capacity of populations to respond evolutionarily to new challenges, such as biotic enemies or changing climate, and consequently increase the risk of local extinction (Barrett & Kohn, 1991; Frankham, 2005). Inbreeding depression is a risk for many rare and threatened species, and can be especially problematic for those that experience rapid population declines and that carry substantial numbers of recessive deleterious alleles (Hedrick & Kalinowski, 2000). Genetic data can help ensure that collections are representative of natural populations and help define management units (Palsbøll et al., 2007) to guide decisions to promote diversity. This can be accomplished effectively and inexpensively through established workflows applying reduced representation sequencing to provide highly informative guidance for conservation management (Rossetto et al., 2021, 2023). Recently it has also become feasible to perform whole genome sequencing (WGS) for specific conservation purposes (Fuentes-Pardo & Ruzzante, 2017; Wright et al., 2020), though this approach remains more expensive than reduced representation sequencing. WGS can offer useful insights about the history of populations, and the frequency of deleterious mutations, which are useful for understanding the risk of inbreeding depression to populations or species.

The long-term goal for species threatened by myrtle rust is to use *ex situ* collections to re-establish self-sustaining populations in the wild. For the most susceptible species, this would require the use of selective breeding, to increase the prevalence of resistance. Resistance breeding has long been carried out in crop species such as wheat, and techniques have advanced with new genomic technologies (Ellis et al., 2014). In eucalypts, it has been possible to breed for resistance to myrtle rust for production forestry (Silva et al., 2013). However, we first need to know whether resistant individuals exist within species that are threatened by myrtle rust, and if so, we need to understand the prevalence and durability of this resistance (under di]erent conditions) (Sniezko et al., 2020). If we can identify resistant individuals using experimental assays, we can begin to contemplate the most appropriate designs for programs of breeding and translocation (Woodcock et al., 2018). These would need to carefully manage the objectives of promoting rust resistant genotypes and phenotypes, while maintaining as much neutral genetic diversity as possible (Bragg et al., 2022; Namkoong, 1991; Dudley et al., 2020). In the absence of endogenous resistance, it might become necessary to contemplate other potential sources. This may include studying the genetic basis of resistance for genetic engineering (Dong & Ronald, 2019) to leverage the immense diversity of plant immune genes (S. H. Chen et al., 2023; Teasdale et al., 2024). Both these approaches have been tested for developing blight-tolerance in the keystone species American chestnut (*Castanea dentata* (Marsh.) Borkh.), though these likely require substantially greater investment and effort than breeding with endogenous resistance (Powell et al., 2019; Newhouse et al., 2014).

In this conservation genomics study, we endeavour to support the long-term management and conservation of *R*. *rubescens* and *R*. *psidioides*. Our primary goal was to characterise the genetic structure and diversity of both species, across their relative distributions, to obtain baseline genomic information to guide efforts in promoting genetic diversity in management activities including to optimise *ex situ* collections. We sought to complement this by making inferences about historical population dynamics and the extent of mutational load, to better understand the risks of inbreeding depression to the managed populations. Finally, we measured rust resistance across *ex situ R*. *rubescens* collections to assess the possible role of breeding as a pathway towards population recovery.

## Materials and methods

### Reduced representation sequencing (DArTseq)

*Rhodamnia rubescens* (*n* = 444) and *Rhodomyrtus psidioides* (*n* = 301) were sampled across their respective known ranges on the east coast of New South Wales and Queensland in Australia (Supplementary Table 1). This sampling of young leaf tissue for the genetic study was performed in conjunction with efforts to collect plant material for vegetative propagation *ex situ*.

Freeze dried leaf tissue samples were sent to Diversity Arrays Technology (DArT) Pty Ltd in Canberra, Australia for DNA extraction and genotyping (DArTseq medium density sequencing) using the documented in-house procedure (Jaccoud et al., 2001). The resulting single nucleotide polymorphism (SNP) data were filtered using approaches and code described in Rossetto et al., 2019 in R v4.1.3 statistical software (R Core Team, 2022). This included the removal of SNPs with a reproducibility score (an index of consistency among technical replicates) of less than 96% and more than 30% missing data.

### Population genomics analysis

Pairwise kinship was estimated using the PLINK method, based on identity-by-descent (Purcell et al., 2007), with SNPrelate v1.17.1 (Zheng et al., 2012) in R. Individuals that are clonal (belonging to the same genet or genetic individual) are expected to have kinship equal to 0.5. However, due to genotyping error, we set a slightly lower threshold value of kinship (0.45–0.5) to infer clonality between plants (Bragg et al., 2020, 2021). Clones detected based on the 0.45 threshold were removed from subsequent analyses.

The R package adegenet v2.1.1 (Jombart, 2008) was used to perform Principal Component Analysis (PCA) on the SNP data to summarise the genetic variation across samples and populations in a small number of uncorrelated variables. This method of PCA derives an ordination based on Euclidean transformed dissimilarity matrix of the data. Population structure was examined by estimating admixture coefficients using sparse Non-Negative Matrix Factorization (sNMF) with the R package LEA v3.16.0 (Frichot & François, 2015) (snmf function with 10 repetitions and *K* values of 2 to 16). Population *F*-statistics and measures of diversity were estimated using the SNPRelate (Zheng et al., 2012) and diveRsity (Keenan et al., 2013) R packages.

### Reference genome sequencing

We generated reference genomes for both *R*. *rubescens* and *R*. *psidioides* and conducted genome-wide analyses of variation. We sampled young leaf tissue from the *ex situ* collections at the Botanic Gardens of Sydney, including *R*. *rubescens* (NCBI BioSample SAMN19602467; collected as NSW1024355 with accession A2017-0001) from tissue culture from PlantBank at the Australian Botanic Garden Mount Annan and *R*. *psidioides* from the living collection at the Royal Botanic Garden Sydney (NCBI BioSample SAMN19602563; collected as NSW1078509 with living collection accession AA16933).

For PacBio HiFi sequencing of *R*. *psidioides*, high-molecular-weight (HMW) genomic DNA (gDNA) was obtained using a sorbitol pre-wash step prior to a CTAB extraction adapted from Inglis et al. (2018). The gDNA was then purified with AMPure XP beads (Beckman Coulter, Brea, CA, USA) using a protocol based on Schalamun et al., 2019 (Lu-Irving & Rutherford, 2021). The quality of the DNA was assessed using Qubit, NanoDrop 1000 (Thermo Fisher Scientific, MA, USA) and TapeStation 2200 System (Agilent, Santa Clara, CA, USA). DNA was sent to the Australian Genome Research Facility (AGRF) where size selection was performed on a BluePippin, using a High Pass Plus Cassette (10-15 kb) (Sage Science, Beverly, MA, USA). Long-read native DNA sequencing was performed on the Pacific Biosciences Sequel II, with an 8M SMRT cell, following the manufacturer’s instructions.

For *R*. *rubescens*, HMW gDNA was obtained using a nuclei isolation (Hilario, 2018) followed by CTAB DNA extraction (Hilario, 2019). Long-read Oxford Nanopore Technologies (ONT) sequencing was performed on the MinION Mk1B using a FLO-MINSP6 (R9.4.1) flow cell with a library prepared with the ligation kit (SQK-LSK109). Basecalling was performed after sequencing with GPU-enabled Guppy v3.2.1. Adapter removal was performed with Porechop v.0.2.4 (Wick et al., 2017) and quality filtering (removal of reads less than 500 bp in length and Q lower than 7) was done with NanoFilt v2.6.0 (De Coster et al., 2018) followed by assessment using FastQC v0.11.8 (Andrews, 2010).

Highly accurate Illumina short-reads were generated for *R*. *rubescens* to polish the ONT assembly. Sequencing was performed at the Ramaciotti Centre for Genomics at the University of New South Wales on an Illumina NovaSeq 6000 with a S2 flow cell with Nextera Flex libraries. Trimmomatic v0.38 was used for adapter removal and trimming with parameters ILLUMINACLIP:${adapters}:2:40:15 LEADING:3 TRAILING:3 SLIDINGWINDOW:4:20 MINLEN:36. Poly-G tails (representing possibly truncated reads) were removed using Bbduk from the BBTools v38.51. Then, a further 15 bases from left and 6 from right of reads were trimmed o] after assessment using FastQC v0.11.8 (Andrews, 2010).

### Reference genome assembly and annotation

The genome assembly workflow varied between the species depending on the sequencing technologies used to generate the data (Supplementary Figure 1). The *R*. *rubescens* genome was assembled using NECAT v0.01 (Y. Chen et al., 2021) followed by tidying with Diploidocus v0.17.0 (S. H. Chen et al., 2022) on cycle mode along with contaminant screening and purging of the myrtle rust pathogen genome (Edwards et al., 2022). Convergence was reached in four rounds. The assembly was long-read polished with Racon v1.4.5 (Vaser et al., 2017) using the parameters -m 8 -x -6 -g -8 -w 500 and medaka v1.0.2 using the r941_min_high model. The short-reads were incorporated by polishing using Pilon v1.23 (Walker et al., 2014) with SNP and indel correction. Scaffolding was done using SSPACE-LongRead v1.1 (Boetzer & Pirovano, 2014) followed by gap-filling using gapFinisher v20190917 (Kammonen et al., 2019) with default parameters. After another round of long-read polishing with Racon v1.4.5 (Vaser et al., 2017) and medaka v1.0.2, we moved forward with a second round of tidying in Diploidocus (default mode). A final polish was performed using Hypo v1.0.3 (Kundu et al., 2019) as it improved the base accuracy of the assembly.

The *R*. *psidioides* HiFi genome was assembled using hifiasm v0.13-r308 (Cheng et al., 2021). The assembly was tidied (haplotig removal and low quality contig trimming) with Diploidocus v0.17.0 (S. H. Chen et al., 2022) on cycle mode (convergence reached in one cycle) followed by vecscreen to screen and filter the genome to remove myrtle rust pathogen contamination, low-quality contigs and putative haplotigs. Gaps in all assemblies were standardised to 100 bp. The *R*. *rubescens* (ToLID drRhoRube1) and *R*. *psidioides* (ToLID drRhoPsid1) assemblies are the first for their species.

Genome completeness was evaluated by BUSCO v3.0.2b (Simão et al., 2015), implementing BLAST+ v2.2.31, Hmmer v3.2.1 and EMBOSS v6.6.0 with the embryophyta_odb9 dataset (*n* = 1,440). For validation and comparison, we also ran BUSCO v5.3.0 (Manni et al., 2021) against the eudicot_odb10 dataset (*n* = 2,326) using MetaEuk (Levy Karin et al., 2020) as the gene predictor. DepthSizer v1.6.2 (S. H. Chen et al., 2022) was used for genome size estimation from the read depth of single-copy orthologues for both species (IndelRatio prediction).

The genomes were annotated using the homology-based gene prediction program GeMoMa v1.7.1 (Keilwagen et al., 2019) with four reference genomes downloaded from NCBI: *Eucalyptus grandis* (GCF_000612305.1; Myburg et al., 2014), *Syzygium oleosum* (GCF_900635055.1; Edwards, unpublished), *Rhodamnia argentea* (GCF_900635035.1; S. H. Chen et al., 2024), and *Arabidopsis thaliana* (TAIR10.1; GCA_000001735.2) (Swarbreck et al., 2007). Annotation completeness was assessed using BUSCO v3.0.2b in proteome mode.

Ribosomal RNA (rRNA) genes were predicted with Barrnap v0.9 (Seemann, 2018) and transfer RNAs (tRNAs) were predicted with tRNAscan-SE v2.05 (Lowe & Chan, 2016), implementing Infernal v1.1.2 (Nawrocki & Eddy, 2013) and the recommended filtering for eukaryotes to form the high-confidence set. A custom repeat library was generated with RepeatModeler v2.0.1 (-engine ncbi) and the genomes were masked with RepeatMasker v4.1.0 (Tarailo-Graovac & Chen, 2009), both with default parameters. The annotation table was generated using the buildSummary.pl RepeatMasker script. Resistance genes were annotated using FindPlantNLRs (S. H. Chen et al., 2023).

### Whole genome resequencing

We chose a set of samples (*n* = 37 for *R*. *rubescens* and *n* = 40 for *R*. *psidioides*) for whole genome resequencing. These samples were chosen with two goals. First, we wanted to target individuals that were distributed broadly across the range of each species. Second, we targeted individuals from *ex situ* collections, to provide molecular information about the specific plants that were available in conservation populations. We also performed whole genome resequencing of a sample of *Rhodamnia sessiflora* and a sample of *Rhodomyrtus longisepala* for use as outgroups.

DNA was extracted from 1-1.5 g of freeze-dried leaf tissue using a modified CTAB method (Doyle & Doyle, 1987) and purified using a Zymo-Spin I-96 Plate and ZR-96 Clean and Concentrator Kit (Zymo Research Corporation, CA, USA) following the protocol of Rutherford et al. (2016). Sequencing was performed at the Ramaciotti Centre for Genomics on an Illumina NovaSeq 6000 with a S2 flow cell with Nextera Flex libraries.

Reads were mapped to the reference genomes of *R*. *rubescens* (drRhoRube1.1) and *R*. *psidioides* (drRhoPsid1.1). AdapterRemoval v2.3.2 (Schubert et al., 2016) was used for adaptor trimming and quality filtering (trimming Phred scores ≤24 with sliding windows of length 8 bp, and output collapsed). Reads were mapped with BWA v0.7.17-r1188 (H. Li, 2013) and processed into BAM files using samtools v1.6 (Danecek et al., 2021). Variants were called using bcftools v1.17 (Danecek et al., 2021) with a mapping quality filter of 30. Sites were filtered for sample depths greater than 6, and genotyping quality greater than 20. Sites with a missingness greater than 6 samples for *R*. *rubescens*, and 7 for *R*. *psidioides*, were also removed. Only biallelic variants were retained, and indels removed. Post-filtering, the coverage of the samples ranged from 9.5x to 31.3x, averaging 20.2x for *R*. *rubescens* and ranged from 17.9x to 29.9x, averaging 24.7x for *R*. *psidioides*. Repetitive regions of both species were masked for SMC++ and diploS/HIC using the RepeatMasker output from the genome annotation. We also masked windows greater than 10 kbp in which repeats were highly prevalent (>50% of bases in repeats).

### Diversity and demographic inference from re-sequencing data

We performed analyses of the whole genome resequencing data to characterize heterogeneity across the genomes in patterns of diversity, and to make inferences about evolutionary history and processes. Both species are experiencing large contractions in their census populations due to the impacts of disease. We therefore sought to understand their demographic histories (in terms of effective population sizes), and variation in the prevalence of putatively deleterious alleles that might influence population management decisions.

We began by estimating levels of genetic diversity (π, *d*_XY_, and *F*_ST_) in moving windows of 100 kb across the genome. To estimate these values, we used scripts available at https://github.com/simonhmartin/genomics_general (accessed 29 November 2023;

Martin & Amos, 2021). We were interested to know if there were regions of the genome with elevated sequence di]erentiation (*d*_XY_, and *F*_ST_), which might be indicative of barriers to gene flow that could be missed with reduced representation sequencing. For these measures, we assigned samples to two populations, from the north and south, in each species (Supplementary Table 1). Negative values of *F*_ST_ were filtered out for plotting.

We inferred demographic histories for both species using SMC++ (Terhorst et al., 2017). This approach infers effective population sizes over historical time points (expressed in terms of generations), using unphased genomic data. Due to uncertainty about the generation length of these species, we only report results in units of generations and do not present a scale with years before present. Parameterisation of generation times were based on estimates of 35 years for *R*. *rubescens* (Threatened Species Scientific Committee, 2020) and 20 years for *R*. *psidioides* (Threatened Species Scientific Committee, 2016). Methods that estimate demographic history are often relatively inaccurate for very recent time points (Sellinger et al., 2021; Terhorst et al., 2017), and we therefore truncated the estimated demographic histories to exclude the first 10,000 generations for the purpose of visualisation.

We sought to understand the deleterious genetic load in these species, and whether it varies in ways that could be useful for managing risks of inbreeding depression. We characterised polymorphic loci in coding regions as synonymous or non-synonymous, and as tolerated or deleterious, using the software package SIFT (Vaser et al., 2016). SIFT uses protein alignments from diverse species to determine whether a specific mutation has been common or rare over evolutionary history, and accordingly infers whether the derived allele is likely to be tolerated or deleterious, respectively. We used SIFT to identify putatively deleterious derived alleles in *R*. *rubescens* and *R*. *psidioides.* We performed analyses to understand whether putatively deleterious alleles varied substantially in their prevalence among individuals, which could be useful in relation to population management. Specifically, we counted the number of predicted deleterious alleles that was observed in each sample. To make sure this index of deleterious ‘load’ was not biased by variation among samples in levels of missing data, we express the number of deleterious alleles in a sample as a fraction of twice the number of non-missing genotype calls (i.e., number of alleles called) at loci having a deleterious allele in the same sample.

Next, we used diploS/HIC v1.0.4 (Kern & Schrider, 2018) to identify regions of the genome that have been subject to selective sweeps. We did this with the hypothesis that strong selection for alleles conferring resistance to myrtle rust might manifest in (soft) sweeps, potentially highlighting genomic regions mediating resistance. Additionally, it has been reported that genomic regions affected by selective sweeps can be enriched with deleterious variants (Schrider & Kern, 2017). If so, this might have implications for identifying regions that mediate inbreeding depression. Using diploS/HIC involves generating simulated genomic sequences evolving under different regimes of selection, including neutral evolution (msprime v1.2.0, Kelleher et al., 2016), hard and soft selective sweeps, or linkage to hard or soft selective sweeps (discoal v0.1.5, Kern & Schrider, 2016). These simulations are otherwise parameterized to reflect the evolution of the relevant population (with effective population size and recombination rate), and their outputs become training data for a machine learning classifier model. This model is then presented with empirical data for genomic variation in windows along the genome and makes predictions for the type of selection that has had a predominant influence on the variation within the region. We were uncertain about the values of some coalescent simulation parameters for these non-model plant species. We therefore ran analyses across plausible ranges of values to check that broad inferences were robust. In light of this, we focus on regions that had large estimated probabilities of having been influenced by soft sweeps. These analyses were limited to genomic scaffolds greater than 500 kbp.

For *R*. *rubescens*, where resistance is more prevalent in *ex situ* collections than in natural populations, we predicted the Gene Ontology (biological processes) of genes in regions that had soft selective sweeps using InterProScan v5.69-101.0 (P. Jones et al., 2014) and the PANTHER v18.0 (Thomas et al., 2022). We tested whether genes involved in particular biological processes were overrepresented in regions that had been subject to soft sweeps (Fisher test implemented via topGO v2.56.0 (Alexa & Rahnenführer, 2009)) relative to genes within all regions of the genome that had diploS/HIC predictions (i.e., excluding genes in highly repetitive regions).

### Myrtle rust resistance assay

We experimentally inoculated individuals from propagated collections of *R*. *rubescens* with the fungal pathogen that causes myrtle rust, and then assessed the development of infection (Sandhu & Park, 2013). We performed these assays for genets from *ex situ* collections that were represented by >3 ramets, to avoid the loss of plants that were needed to maintain diversity in the collection. We also performed assays for seedlings raised from seeds produced by the collections. We did not perform assays for individuals in the *R*. *psidioides* collection, as fewer ramets were available for each genet of this species. Most plants in these *ex situ* collections are routinely treated with fungicide, including 56 that were assayed in this study. These treatments were suspended at least four weeks before the assay to ensure there was no residual effect (Carnegie et al., 2016). A remaining 241 seedlings that were assayed were never treated with fungicide. Plants were inoculated with the urediniospores from the pathogen *A*. *psidii* using a protocol described in Sandhu & Park (2013). Briefly, urediniospores were multiplied on a susceptible *Syzygium jambos* L. (Alson) host. Inoculations were performed using a fine mist spray, applied at a concentration of 2.0 mg urediniospores suspended in 1.0 mL of light mineral oil (Univar Solvent L Naphtha 100, Univar Australia Pty. Ltd.). Post inoculation, plants were placed into a 20 °C, dark incubation chamber under constant misting for 24 h. Plants were then maintained at 22 ± 2 °C in a naturally lit microclimate room prior to disease assessment at 14-day post-inoculation and scored for manifestations of infection and host responses according to Supplementary Table 2 and also assigned a binary score of 0 (resistant) or 1 (susceptible). Rust resistance assays were carried out at the Plant Breeding Institute (PBI) Cobbitty at the University of Sydney. Fourteen assayed plants were planted out in the field in northern New South Wales, in a region conducive to the development of rust infection, for further observations.

## Results

### Reference genomes for two critically endangered Myrtaceae

For *R*. *rubescens*, sequencing yielded 24.1 Gbp of ONT MinION data and 36.5 Gbp of Illumina reads. For *R*. *psidioides,* PacBio HiFi sequencing yielded a total of 19.2 Gbp of data. Sequencing output for each plant and platform is available in Supplementary Table 3.

The genome size of the studied species had not been previously measured using flow cytometry. For *R*. *rubescens*, DepthSizer predicted a genome size of 352 Mbp (Supplementary Table 4) whilst the drRhoRube1.1 assembly was slightly larger at 354 Mbp (174 scaffolds with N50 = 3.7 Mbp). The drRhoPsid1.1 assembly had a larger size at 931 Mbp and while there was a higher number of scaffolds at 828, it was more contiguous with N50 = 5.7 Mbp (Table 1). DepthSizer predicted a genome size of 860 Mbp (Supplementary Table 4).

**Table 1.**
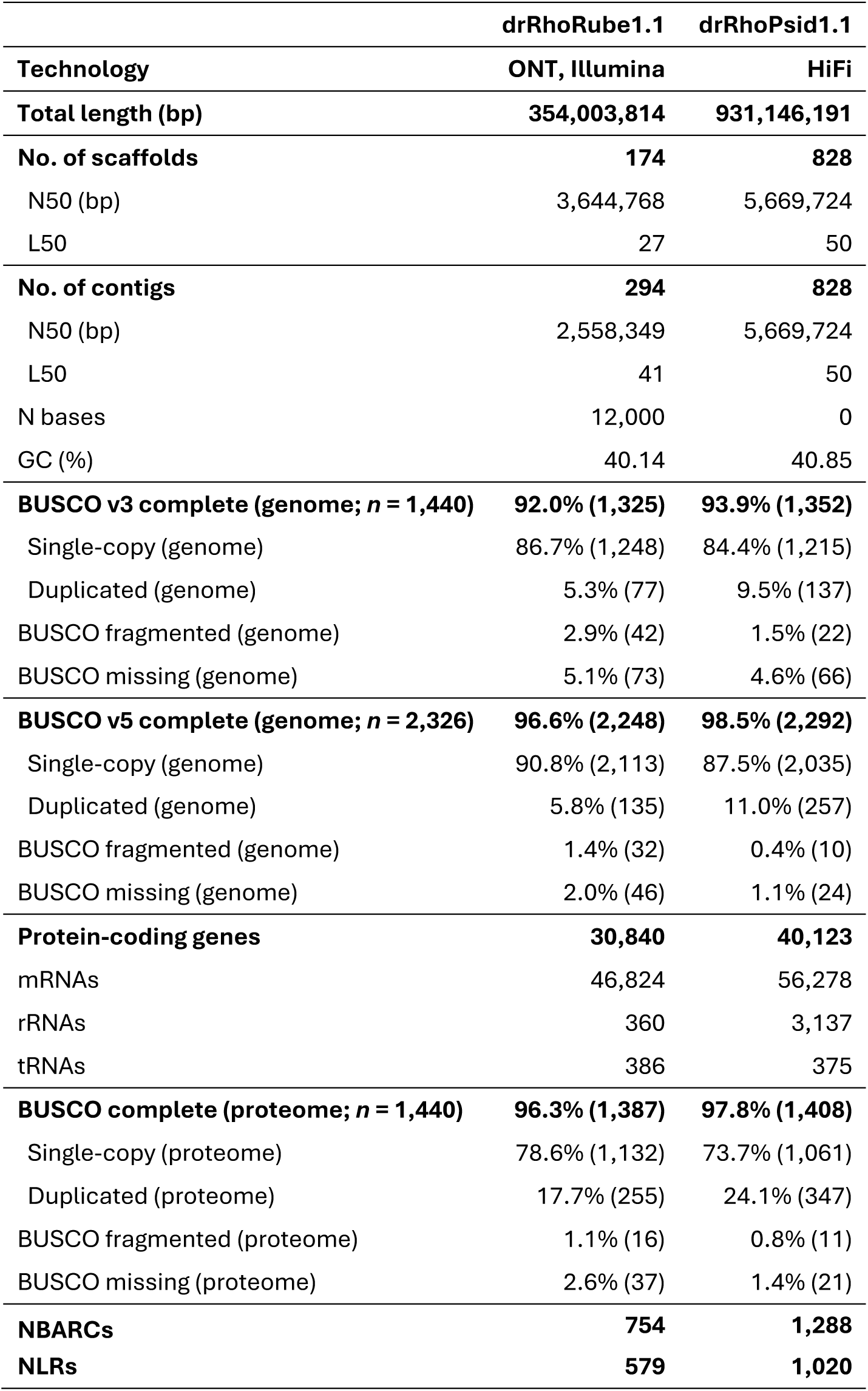
Genome assembly and annotation statistics for *Rhodamnia rubescens* (drRhoRube1.1) and *Rhodomyrtus psidioides* (drRhoPsid1.1).

BUSCO v5 completeness was high for both plant genomes, being 96.6% for the drRhoRube1.1 assembly and 98.5% for the drRhoPsid1.1 assembly, with the vast majority of genes being complete and single-copy. For drRhoRube1.1, there were 30,840 genes, 360 rRNAs, and 386 tRNAs (Table 1). The genome had a repeat content of 47.5% (Supplementary Table 5). The annotation of drRhoPsid1.1 revealed 40,123 protein coding genes, with a high number of rRNAs at 3,137 but a similar number of tRNAs to drRhoRube1.1 at 375. The genome was also highly repetitive (75.1%). The proteomes had a comparable level of completeness to the genomes. For drRhoRube1.1, there were 754 putative NBARC-containing genes and 579 NLR genes. For drRhoPsid1.1, there were 1,288 NBARCs and 1,020 NLRs (Table 1).

### Population genetic diversity and structure

DArTseq sequencing of *R*. *rubescens* samples collected from the wild (*n* = 444 across 72 localities) yielded 29,997 SNPs which were quality filtered to 16,834 SNPs.

A minor allele frequency (MAF) filter of 0.05 was applied to each SNP dataset to obtain common variants for the kinship analyses. For *R*. *rubescens*, this reduced the dataset to 4,608 SNPs. The analysis detected 148 samples were ramets representing 50 genets (Supplementary Figure 2 and Supplementary Table 6). All samples inferred to be clonal were collected from the same locality. The genet with the most ramets (7) was located at Black Head and another large genet (6 ramets) was detected at Diamond Head within Crowdy Bay National Park.

*Rhodomyrtus psidioides* (*n* = 301 across 45 localities) yielded 22,338 SNPs. This was reduced to 14,008 SNPs after quality filtering, and 5,706 common SNPs for the kinship analyses. The analysis detected 183 out of the 301 samples were clonal. There were 67 genets comprising of multiple ramets, with most of these genets having 2 ramets (Supplementary Figure 3 and Supplementary Table 7). Notably, the largest genets, each of 7 ramets, were located at Red Rock and Wamberal.

Genetic variation is primarily distributed across a latitudinal gradient for both species along the east coast of Australia (Figure 1). In *R*. *rubescens*, the samples from localities near the Sydney area appear to be outliers in the PCA, suggesting they are planted. In *R*. *psidioides*, the Wahroonga and Pearl Beach populations were similarly outliers compared to others, suggesting that these were planted populations. Admixture analysis (sNMF) suggests that genetic variation is continuous in *R*. *rubescens* (i.e., not discretely structured). The analysis for *K* values from 2 to 16, suggested a large optimal value (*K* = 13). For *K* = 2, a gradient was apparent across latitudes (Supplementary Figure 4A). However, structure is more pronounced in *R*. *psidioides* (*K* = 3) with similar gradient across latitude (Supplementary Figure 4B). For *R*. *rubescens,* values of observed heterozygosity (*H*_O_) ranged from 0.12 to 0.24 and expected heterozygosity (*H*_E_) ranged from 0.08 to 0.19. The inbreeding coe]icient (*F*_IS_) was negative for *R*. *rubescens* sites due to the small population size and therefore unreliable. For *R*. *psidioides*, we present *F*-statistics according to the three admixture groups. *H*_O_ ranged from 0.16 to 0.19 while *H*_E_ ranged from 0.28 to 0.30, indicating a deficiency of heterozygotes. Inbreeding was detected in *R*. *psidioides* with *F*_IS_ ranging from 0.34 to 0.39 (Supplementary Table 8 and Supplementary Table 9).

**Figure 1.**
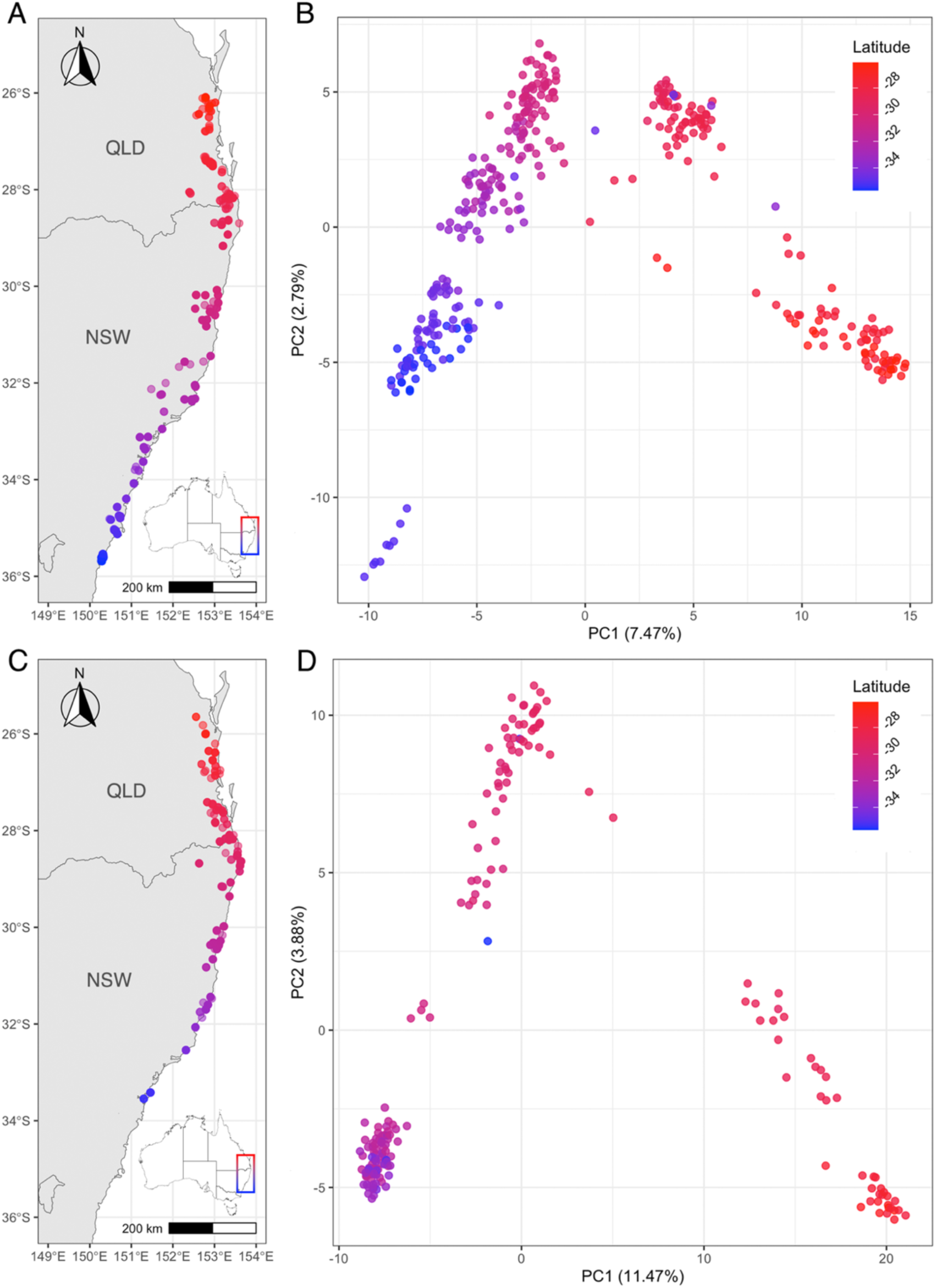
Distribution of genetic variability along a latitudinal gradient for two previously common but now Critically Endangered species due to high susceptibility to myrtle rust. Comprehensively sampled distribution across eastern Australia and PCA (PC1 axis vs PC2 axis) based on SNP data for **A–B** *Rhodamnia rubescens and* **C–D** *Rhodomyrtus psidioides,* with points coloured by latitude.

### Whole genome resequencing

We generated whole genome resequencing data for 37 individuals of *R*. *rubescens* and 40 individuals of *R*. *psidioides.* A number of these individuals turned out to be related or clones, and we present analyses of 25 (*R*. *rubescens*) and 37 (*R*. *psidioides*) unrelated individuals (Supplementary Table 1). Whole genome resequencing of *R*. *rubescens* and *R*. *psidioides* yielded 15.6 and 13.6 million SNPs, respectively. The two species exhibited similar levels of genome-wide diversity and differentiation (Supplementary Figure 5). For *R*. *rubescens*, mean nucleotide diversity (ρε) was 0.194 (range: 0.061 to 0.278), and for *R*. *psidioides* mean nucleotide diversity was 0.206 (range: 0.031 to 0.315). Both species exhibited low levels of genetic differentiation (*F*_ST_) between regions, with *R*. *rubescens* averaging 0.021 and *R*. *psidioides* averaging 0.024. We did not identify any regions of the genome that were outliers with large values of *d*_xy_ or *F*_ST_.

### Historical demography

*Rhodamnia rubescens* and *R*. *psidioides* both had large estimated effective population sizes (>100,000) for most of the last 400,000 generations. Both species also exhibit trajectories of mild contraction from approximately 250,000 generations ago (7.5 Mya, if generation time is 30 years; Figure 2).

**Figure 2.**
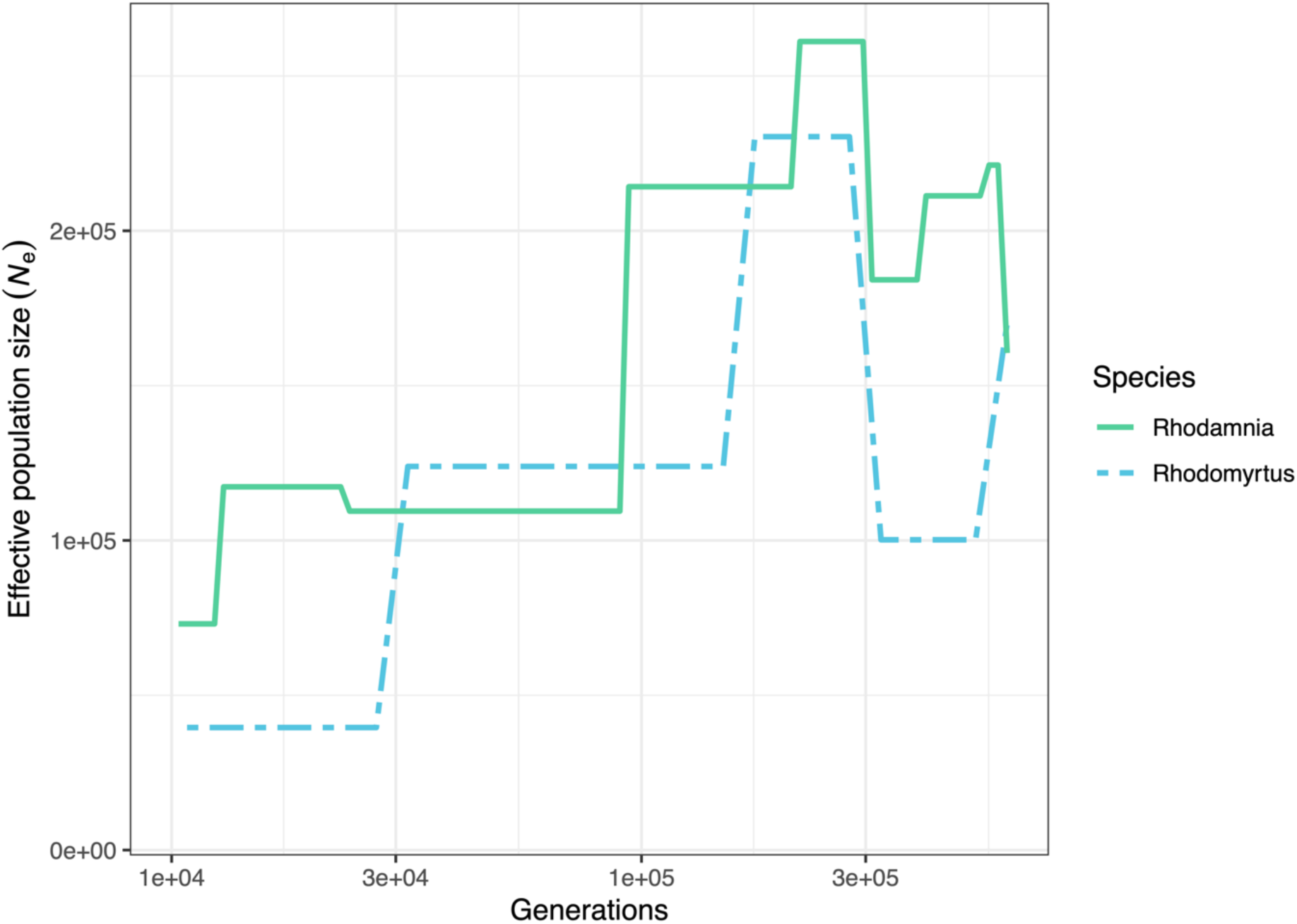
Historical demography inferred using SMC++ for *Rhodamnia rubescens* (green solid line) and *Rhodomyrtus psidioides* (blue dashed line). Inferred effective population size (*N*_e_) is shown as a function of generations before present (note log axis scaling).

### Selection and deleterious load

In *R*. *rubescens*, 939,083 SNPs were identified in protein sequences by SIFT and classified as either deleterious (14.9%) or tolerated. In *R*. *psidioides*, 568,310 deleterious or tolerated SNPs were identified in protein sequences, and a similar fraction (15.8%) were deleterious. Among SNP loci, frequencies of derived deleterious alleles tended to be lower than frequencies of derived tolerated alleles in both species (Figure 3A–B). Overall, there was relatively little variation among *R*. *rubescens* individuals in the frequencies of deleterious alleles (per called alleles at sites with a deleterious allele), except for one outlier sample, which had a substantially lower frequency of deleterious alleles (Figure 3C). In *R*. *psidioides*, there was a tendency for samples from the north of the range to have greater frequencies of deleterious alleles than individuals from the south (*t*_18.4_ = -10.18, *p* = 2.8 x 10^-9^), though the difference in means between the groups was relatively modest (0.155 vs 0.167).

**Figure 3.**
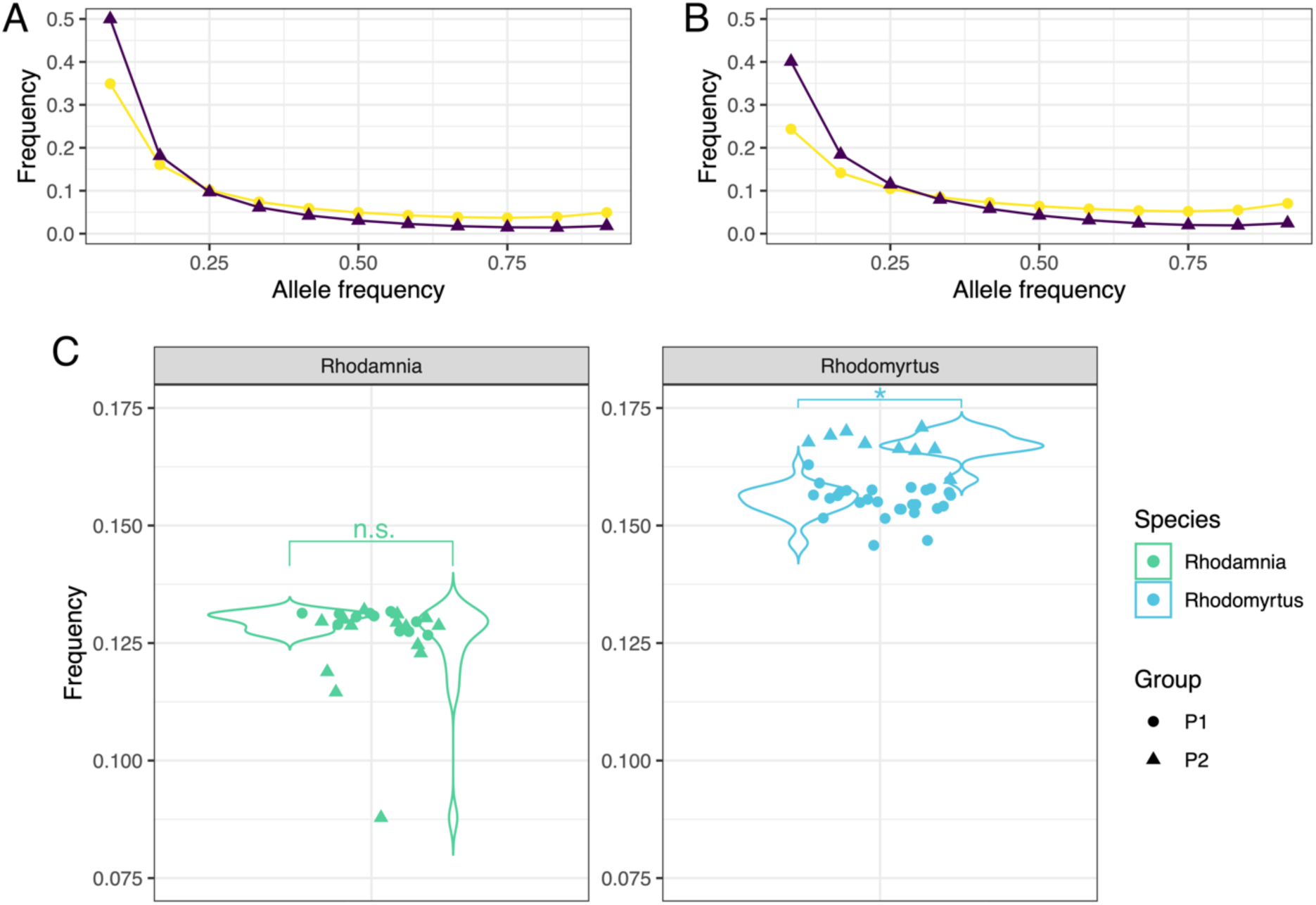
Population frequencies for tolerated (yellow circles) and deleterious (purple triangles) derived alleles from SIFT analysis for **A** *Rhodamnia rubescens* (*n* = 25) and **B** *Rhodomyrtus psidioides* (*n* = 37). **C** Individual levels of deleterious alleles.

For *R*. *rubescens*, 6.8% of genome regions were predicted by diploS/HIC to have been subject to soft selective sweeps with greater than 90% probability. This declined to 0.7% of regions for soft sweeps with predicted probabilities greater than 99.9% (Supplementary Figure 6). For *R*. *psidioides*, 2.5% and 0.05% of regions were predicted to have had soft sweeps with probabilities exceeding 90% and 99.9%, respectively. In *R. rubescens*, 10 Gene Ontology categories were overrepresented (Fisher test *P*<0.01) in regions that were predicted (with probability >90%) to have had soft selective sweeps (Supplementary Table 10). This included a group of genes involved in broad responses to stress (GO:0033554). Using Gene Ontologies, we also identified 2 genes involved in plant hypersensitive responses (GO:0009626), and 3 genes involved in immune responses (GO:0006955), that occurred in regions that had soft selective sweeps. Among the NLR and NBARC genes annotated in the *R. rubescens* genome, one NBARC gene (g3063) occurred in a region that was subject to a soft selective sweep.

In *R*. *rubescens*, regions of the genome that were linked to hard and soft selective sweeps were modestly enriched with deleterious SNPs relative to neutral regions. The deleterious to tolerated ratio was 0.157, 0.17, and 0.164 for neutral, linked hard, and linked soft regions, respectively (Fisher test *P* = 0.015 for linked hard regions and *P* = 0.0008 linked soft regions). Regions with soft or hard sweeps were not significantly enriched with deleterious SNPs relative to neutral regions. In *R*. *psidioides*, we observed similar patterns, with regions linked to hard and soft selective sweeps having elevated numbers of deleterious SNP loci. The deleterious to tolerated ratio was 0.17, 0.23, and 0.19 for neutral, linked hard, and linked soft regions, respectively (Fisher test *P* = 0.02 for linked hard regions and *P* = 0.0003 for linked soft regions).

### *Rhodamnia rubescens* exhibits variation in myrtle rust resistance

There was a spectrum of responses to myrtle rust observed in *R*. *rubescens*, with plants varying from highly susceptible to highly resistant (Supplementary Figure 7). Of the 297 assayed plants, 163 (55%) displayed resistance and 134 (45%) were susceptible (Supplementary Table 11). The assayed plants were from two *ex situ* collections, held at the Australian Botanic Garden Mount Annan (*n* = 259) and Booderee Botanic Gardens, and included individuals grown from cuttings and from seeds produced in the *ex situ* collections. Seedlings (*n* = 241) from 22 propagated plants held at the Australian Botanic Garden Mount Annan exhibited substantial variation in levels of resistance. For accessions with more than 10 assayed seedlings, the proportion of resistant seedlings varied from 9% to 86%. Collectively, where the phenotype of the mother plant was resistant (10 families), seedlings were on average 67% resistant. Where the phenotype of the mother was susceptible (2 families), the seedlings were on average 51% resistant (Supplementary Table 12). Following the assays, a total of 14 plants that were found to be resistant were planted and monitored for two years at sites in northern New South Wales, Australia. Of these, 10 plants (71%) developed substantial infection (severe dieback or death); only 2 plants were unaffected by rust with no dieback observed (Supplementary Table 11).

## Discussion

Emerging fungal pathogens can impact host populations rapidly. To conserve heavily impacted host species, conservation practitioners must act quickly to secure populations to limit the loss of genetic diversity, while searching for ways to establish self-sustaining, resistant, translocated populations (Woodcock et al., 2018). The disease myrtle rust has caused very large declines in several Myrtaceae species in Australia in a single generation (Carnegie et al., 2016; Fensham et al., 2020, 2021). For two species that are among the most severely affected, *R*. *rubescens* and *R*. *psidioides*, this study has generated information that can guide management from the level of broad strategic planning (e.g. highlighting the potential importance of inbreeding), to specific aspects of implementation (e.g. identifying putatively resistant individuals). These approaches are applicable to other species facing threats from novel pathogens, and other threats that lead to very rapid losses of individuals and diversity.

The overarching strategy for conserving *R*. *rubescens* and *R*. *psidioides* is to protect and preserve as much of the remaining diversity as possible, using *ex situ* collections and other methods of germplasm storage (Sommerville et al., 2019), while plans are devised for breeding to improve disease resistance (Makinson et al., 2020). This study underpins the notion that until recently, both species have been genetically diverse, with large effective population sizes, and substantial gene flow across the landscape. This raises the possibility that these species might be at risk of inbreeding depression, following a large population contraction (Kyriazis et al., 2021). Our observations are consistent with this, identifying overrepresentation of deleterious alleles at low frequencies, and linked to past selective sweeps, consistent with previous observations (Schrider & Kern, 2017).

We did not observe variation among individuals in levels of deleterious alleles that might be usefully exploited in breeding programs (Speak et al., 2024). However, we did observe slightly elevated frequencies of deleterious alleles in *R. psidioides* individuals from northern NSW, and this might be weighed in relevant population management decisions. More broadly, our observations highlight that both *R. rubescens* and *R. psidioides* face risks from inbreeding depression, and management actions should accordingly place emphasis on the avoidance of inbreeding.

For both species, levels of genetic connectivity across the landscape were relatively high, with little evidence of strong differentiation between putative groups, or conspicuous barriers to gene flow in the form of islands of divergence. In sum, these observations do not point to any problems with mixing germplasm from widespread collection sites, from the perspective of genetic compatibility. For these species which are undergoing massive contractions in effective population sizes and with limited numbers of unrelated individuals in each collection, it might be advantageous to promote mixing. We do note, however, that these observations do not address other important considerations, such as the possibility of local adaptation to prevailing environmental conditions by populations in different parts of the range.

This study also highlights the potential feasibility of breeding for rust resistance in *R*. *rubescens*. A small number of plants showed resistance to myrtle rust both in experimental assays and when planted in a region with a climate that is conducive to infection. If this resistance is heritable, stable under the environmental conditions prevailing in potential translocation sites, and lasts into reproductive maturity, it will provide pathways towards breeding populations with durable disease resistance (Sniezko et al., 2020). We note that other plants developed infection and exhibited dieback when planted in the trial, despite appearing resistant in experiments (Supplementary Table 11). There are several possible explanations for this, including the effects of residual fungicide, nutrition, stressors, or the growth stage of leaves (J. D. G. Jones & Dangl, 2006; Carnegie et al., 2016; Beresford et al., 2020; Develey-Rivière & Galiana, 2007). Taken together, these results highlight that breeding programs for conservation might benefit from a combination of experimental assays to quickly identify and exclude highly susceptible progeny, followed by field trials and monitoring of individuals that are putatively resistant. Future studies may focus on establishing the genetic basis of resistance to myrtle rust and identifying candidate loci to inform targeted screening and breeding programs (Periyannan et al., 2017; Stocks et al., 2019; Weiss et al., 2020; Yong et al., 2021). This study adds to the high quality genomic resources for the Myrtaceae family (F. Li et al., 2023), particularly for myrtle rust impacted species, which will be essential to the search for associated markers and broader aspects of conservation management (Brandies et al., 2019; Hogg et al., 2022). For *R*. *psidioides*, it is not yet clear whether there is a pathway towards breeding using resistance that occurs within the species, or if it will be necessary to search for alternatives, either involving introgression or biotechnological approaches for introducing resistance to populations (Woodcock et al., 2018).

A rapid and coordinated response across the distribution of myrtle rust impacted species is essential for securing species such as *R*. *rubescens* and *R*. *psidioides* (Makinson et al., 2020). This conservation framework may also be extended to other susceptible species. Conservation actions should be guided by genomic information generated as part of this study so that genetic diversity and resilience can be promoted, and resources are efficiently used to enhance the long-term survival of myrtle rust impacted species.

## Acknowledgements

We acknowledge the Traditional Owners and Custodians of the lands on which this research was conducted. We acknowledge the many collectors with relevant permits who assisted with field work and property owners for access to plants. In particular, we thank Elliot Bowerman at the Sunshine Coast Council and Nick Swanson at Logan City Council for assistance in the field. We thank those who responded to the request on the NSW Native Plant Identification Facebook group regarding *Rhodamnia rubescens* sightings, in particular, Peter Woodward, Jill Albrecht, and Lily Mickaill. We thank Richard Dimon and Kit King for processing samples and metadata at the Research Centre for Ecosystem Resilience at the Botanic Gardens of Sydney. We thank Elena Hilario and Benjamin Schwessinger for assistance with HMW DNA extraction and MinION sequencing during a workshop at the Australian National University.

## Data availability

The *Rhodamnia rubescens* genome (drRhoRube1.1), raw data (Nanopore long-reads and Illumina reads), and whole genome sequencing data were deposited to NCBI under BioProject PRJNA735913. The *Rhodomyrtus psidioides* genome (drRhoPsid1.1), raw data (PacBio HiFi), and whole genome sequencing data were deposited to NCBI under BioProject PRJNA735915. DArTseq data and genome annotation files were deposited to the CSIRO Data Access Portal (https://doi.org/10.25919/j15r-tt10).

## Benefit-sharing

All collaborators are included as co-authors. The research addresses the conservation of plant species being studied. All data have been shared with the broader public via appropriate biological databases.

## Author contributions

The eleven middle authors are (reverse) alphabetised by last name and considered equal. The first author and three senior authors conceptualised and coordinated the project. Stephanie Chen performed the DArTseq data analysis with guidance from Samantha Yap and Jason Bragg. Stephanie Chen extracted the HMW DNA for the reference genomes and Ashley Jones performed the Nanopore sequencing. Samantha Yap performed the extractions for the resequencing. Stephanie Chen, Richard Edwards, and Jason Bragg assembled and annotated the reference genomes. Jason Bragg, Karina Guo, and Stephanie Chen performed the analyses of the resequencing data. Craig Stehn led the field work in New South Wales and was assisted by Joel Cohen, Stephanie Chen, Jason Bragg, Samantha Yap, and Maurizio Rossetto. Fiona Giblin led the field work in Queensland and was assisted by Tracey Menzies, Geoff Pegg, and Craig Stehn. Veronica Viler maintained and curated the *ex situ* collection at Australian Botanic Garden Mount Annan and Julie Percival maintained and curated the *ex situ* collection at Booderee Botanic Gardens. Karanjeet Sandhu performed the rust assays. Stephanie Chen and Jason Bragg wrote the manuscript. All authors read and approved the final manuscript.

